# Nischarin negatively impacts ShcD-mediated tumor cell migration

**DOI:** 10.1101/2023.07.10.547766

**Authors:** Rayan A. Hago, Sook P. Wong, Mahmood Y. Hachim, Ibrahim Y. Hachim, Maha Saber-Ayad, Sally A Prigent, Samrein B.M. Ahmed

## Abstract

ShcD was previously found to promote cell motility in melanoma cells. Screening of a yeast two hybrid mouse embryo cDNA library identified Nischarin, a negative regulator to cell motility, as an interacting partner to the ShcD-CH2 domain. Therefore, we aimed to investigate the interaction between Nischarin and ShcD in mammalian cells and to determine their functional impact on cell migration. The Nischarin/ShcD interaction was confirmed by transfection and co-immunoprecipitation assays using full-length constructs in HEK293, MCF7 and MM253 cell lines. Deletion of the first 93 amino acids of ShcD abrogated the interaction indicating the importance of these residues for Nischarin binding. Co-expression of Nischarin and ShcD demonstrated an inhibitory effect on the levels of phospho-ERK and phospho-LIMK. In support of this, Nischarin was found to block the migratory activities of ShcD. A brief in silico analysis of publicly available breast cancer patient data was performed to elucidate the effect of Nischarin/ShcD co-expression on the patients’ overall survival. Patients with high expression of both proteins had better survival than those with only ShcD overexpression. Our results reveal that the novel protein Nischarin is an interacting partner to ShcD. In addition, we report that the tumour suppressive abilities of Nischarin can overcome ShcD-mediated cell migration when both proteins are concomitantly expressed.

*This abstract was presented in the National Cancer Research Institute (NCRI)-2019

## Introduction

Adaptor proteins facilitate protein-protein interactions, which are responsible for intracellular signal transduction and protein organisation. The Src homology and collagen (Shc) family consists of a group of adaptor proteins that are evolutionarily conserved and share functional and structural features (1). All Shc proteins contain an amino-terminal phosphotyrosine-binding (PTB) domain, a linker collagen homology 1 (CH1) domain and a carboxy-terminal Src homology 2 (SH2) domain. An additional amino-terminal collagen homology region (CH2) is encoded in the longest Shc protein transcripts (1, 2). The Shc family consists of four members encoded by four distinct genes: Shc/Shc1/ShcA, Sli/Shc2/ShcB, Rai/Shc3/ShcC and RaLP/Shc4/ShcD (1, 3, 4). Broadly, Shc proteins mediate several intracellular signalling cascades and are implicated in cell proliferation, cell differentiation, tumorigenesis, and cell survival and migration (5–10). Despite their similar structure, differing roles have been ascribed to the individual Shc family members and they are differentially expressed in tissues (1).

The reported novel roles of ShcD include triggering melanoma invasion and neuromuscular junction maturation as well as functioning in mouse embryonic development (3, 4, 11). Recently, Aladowicz et al reported an interaction between ShcD and Dock4 which impacted Rac1 signalling and cell motility in melanoma cells (12). Despite the reported intracellular roles of ShcD, the exact molecular mechanisms by which ShcD mediates these biological responses are not fully revealed.

One approach to elucidate the role of ShcD in intracellular signalling is to identify and study its interacting partners. ShcD was found to interact with various tyrosine kinase receptors through its PTB and SH2 domains (13, 14). A SPOT peptide array was used to identify binding partners for different PTB domains from various adaptors using a human library of NPXY motifs, which revealed that the ShcD-PTB domain interacted with TRKB/C, Met, Ret, IGF1R, ErbB2/4 and VEGFR3. Unlike the PTB domain of other Shc proteins, the ShcD-PTB domain binds the muscle-specific kinase receptor (MUSK) (13).

Herein, we aimed to identify ShcD-interacting proteins that might further elucidate the novel role of ShcD in cell migration. We focussed on the N-terminal CH2 domain of ShcD as there have been no proteins previously identified to bind to this region. Since this region is poorly conserved between Shc proteins it may be responsible for the divergent functions of different family members. In the case of ShcA, different splice variants with the presence (p66Shc) or absence (p52Shc) of the CH2 domain have very different functions; the former being associated with stress induced apoptosis, with the latter having a well-established role in tumour cell proliferation (15–17). The ShcD-CH2 domain was used to screen a mouse embryo yeast two hybrid cDNA library. Nischarin, a novel protein and a negative regulator of cell motility (18, 19), was identified as a binding partner. The functional consequence of this interaction was explored in MCF7 and MM253 cell lines. The rationale of using these two cell lines is that Nischarin was found to have a tumour-suppressive effect in breast cancer (20), while ShcD was reported to act as an oncogene in melanoma cells (3). Moreover, both proteins are expressed in melanoma and breast cancer as it is demonstrated by Human Protein Atlas (Figure 1SA & 1SB) (21).

## Materials and Methods

### Reagents and antibodies

All reagents used for cell line maintenance were supplied by Sigma-Aldrich, UK. Reagents that were used in the western blot technique and other molecular biology techniques were supplied by Thermo Fisher Scientific, USA and Sigma-Aldrich, UK. Anti-FLAG (F1804-Sigma-Aldrich, UK), anti-GFP (sc9996-Santa Cruz, USA), anti-phospho Erk1/2 (cst4370S-Cell Signaling, USA), anti-Erk1/2 (cst4695-Cell Signaling, USA), anti-Nischarin (ab204588-Abcam, UK), anti-phospho-LIMK (3841S-Cell Signaling, USA), and anti-LIMK (3842S-Cell Signaling, USA) antibodies were used.

### Cell lines

The breast cancer cell line MCF7 was kindly gifted by Prof. Rafat Alawaady (University of Sharjah, UAE). A melanoma cell line (MM253) was purchased from the European Collection of Authenticated Cell Cultures (ECACC)/UK.

### Constructs

FLAG-tagged ShcD constructs were generated by the Protex facility (Department of Molecular and Cell Biology, University of Leicester, UK). The full length ShcD sequence provided at accession NM_203349.3. The vector name used for cloning full length and truncated human ShcD is pLEICS-12; available at https://le.ac.uk/mcb/facilities-and-technologies/protex/available-vectors/mamalian-expression. All the tags were linked at the amino terminus of the ShcD sequence. The sequences of the constructs were verified by DNA sequencing at the PNACL facility (University of Leicester, UK). The GFP-Nischarin (mouse) expression vector was a generous gift from S. Alahari (Rockefeller University, New York) and the vector was described in Alahari et al., 2000 (19). FLAG negative, empty vector (EV), (CV016) and GFP vector (CV026) were purchased from Sinobiological, China. The DNA sequence corresponding to the ShcD- CH2 domain (amino acid 1-179) was cloned into the SalI site of the vector pBTM116- PDGFR (22) for yeast two hybrid library screening.

### Yeast culture and library screen

Yeast two hybrid screening was performed using the ShcD-CH2 domain fused to LexA DNA binding domain, and a mouse embryo library constructed in the pVP16 vector which encodes the VP16 activation domain (23). The L40 yeast strain (MATa trp1 leu2 his3 LYS2::lexA-HIS3 URA3 ::lexA-LacZ) was used (24). Yeast culture, transformation and library screen were conducted as described previously (25). Yeast colonies growing on selective plates lacking tryptophan, leucine and histidine were tested for their ability to activate the β-galactosidase reporter using filter assays with X-gal substrate (26). Plasmids were isolated from positive colonies and retransformed into yeast with a negative control plasmid (lamin-pBTM116) to test for non-specific binding. Plasmid inserts were sequenced by PNACL, University of Leicester.

### Cell lines maintenance and transfection

MCF7 and MM253 cell lines were cultured in RPMI-1640 medium (R8758-Sigma Aldrich, UK) supplemented with 10% foetal bovine serum (P9665-Sigma Aldrich, UK) and 1% Penicillin-Streptomycin solution (P4333, Sigma Aldrich, UK). MM253 cells medium contained 25 mM HEPES in addition to the previously stated constituents. The cells were incubated in a humidified incubator connected to 5% CO_2_ and the temperature was adjusted to 37 _JC. The cells were maintained under maximum possible aseptic conditions. Transient transfections were performed using Viafect transfection reagent (E4981-Promega, USA) following the manufacturer’s guidelines.

### Coimmunoprecipitation

The transfected cells were lysed with Triton-based lysis buffer, supplemented with 1mM PMSF, 1 mM Na_3_VO_4_, 50 mM NaF and protease inhibitor cocktail. After removing the insoluble fraction of the cell lysate, the supernatant was transferred to G Sepharose beads (Sigma-Aldrich; P3296) coupled with 2 µg of anti-FLAG antibody. The lysates were left tumbling with the antibody-coupled beads overnight. The beads were then washed, and sample buffer containing DTT was added to detach the proteins from the beads. In the case of GFP-Nischarin immunoprecipitation, GFP-Trap agarose (Chromotek, Germany) was used following the manufacturer’s guidelines.

### Western blotting

Soluble proteins obtained from cell lysates or from the co-immunoprecipitation experiments were resolved on SDS-PAGE gels, and the proteins were transferred to PVDF membranes. After the membrane was blocked, diluted primary antibodies were incubated with the membrane overnight at 4°C. The secondary antibodies use was based on the primary antibody species. Enhanced chemiluminescence was employed to detect the protein of interest (34580, Thermo Scientific™, USA). The bands were then visualized with a Chemidoc machine (BioRad). When re-probing was required, stripping buffer (Restore™ PLUS Western Blot Stripping Buffer; 46430) was used. Band intensities were quantified by ImageJ software.

### Immunofluorescence

The cells were seeded on sterile cover slips and fixed with 3.7% formaldehyde for 20 min at RT. Cells were permeabilised with 0.1% Triton X-100 in PBS for 10 min followed by a blocking step using 3% BSA in PBS. Primary antibody was added for 1 hr at RT. After the primary antibody was washed off, secondary antibody was added for 1 hr at RT. The nuclei were then stained with DAPI contained in the mounting medium (Abcam; ab104139) for 5-10 min. The coverslips were then sealed. The cells were visualised using a fluorescence microscope (Olympus, BX51TF) and CellSens standard software. For ShcD and Nischarin co-localization, Alexa Fluor 495 (red) was used to visualize FLAG-ShcD. The overlay was performed using ImageJ-win64 software.

### Wound healing assay

The cells were cultured in a 6-well plate until they reached 80% confluence; the cells were transfected. After 24 hrs, the wounds were made to the confluent monolayer of cultured cells. The cells were washed once with PBS, and phenol red-free compatible medium was added. Wound imaging was performed using an Optika microscope, Optika camera (XDS-2) and a 10x objective. The images were taken at 0 hrs and 24 hrs using Optika software and the analysis was performed using ImageJ-win64 software.

### Transwell assays

For the transwell assays, 1.25x10^5^ cells were added to the upper chamber of collagen-coated Boyden chamber containing a membrane with 8-µm pores. The cells were resuspended in 0.1% FBS-containing medium, while the lower chamber contained 10% FBS-containing medium. The cells were incubated for 24 hrs. The cells at the bottom surface of the membrane were stained with crystal violet (1:20) and solubilised with 4% SDS. Separate tubes contained 1.25x10^5^ from each condition were pelleted and stained with crystal violet and dissolved by SDS representing the total number of cells. The colorimetric reading was taken at 570 nm. The percentage of the migrated cells was calculated by dividing the colorimetric reading of the migrated cells by the colorimetric reading of the total number of cells.

### Studying the association between the mRNA expression of Nischarin and ShcD in samples from a large cohort of breast cancer patients via bioinformatics

The bc-GenExMiner 4.2 database tool (27, 28) was used to investigate the correlation between mRNA expression levels of Nischarin and Shc4 and tumour grade as well as molecular subtypes in a large patient cohort. The Kaplan-Meier (KM) plotter database (29) was used to evaluate the association between Nischarin and SHC4 expression levels and patient outcomes presented as relapse-free survival (RFS) and overall survival (OS).

This database uses automatic beforehand method that allows classification of samples according to genes median expression through auto select best cut-off value. Patients that showed gene expression levels below the median were classified as low and patients express levels above the median were classified as high expressors.

Using the multiple genes tools in the Kaplan-Meier (KM) plotter database, we were able to generate a gene signature composed of the mean expression levels of both genes to evaluate the effect of combined expression of both genes in patients. Next, the association between this gene signature and patient outcomes, presented as relapse free survival (RFS) and overall survival (OS), was evaluated.

### Statistical analysis

One-way Anova test was employed to determine the statistical significance between the different conditions. When Anova test showed significance, multiple comparison was conducted between the different conditions. The p-value was considered significant at <0.05. Error bars represent the standard error of the mean (SEM). GraphPad prism 7.04 was used for the data analysis and data presentation. Band intensity in western blotting was measured using ImageJ-win64 software (30).

## Results

### The ShcD-CH2 domain interacts with the invasion suppressor, Nischarin

Two positive colonies were identified though the yeast two hybrid screen. Sequencing of the plasmids isolated from these colonies revealed two overlapping sequences encoding part of the Nischarin protein, designated 6.1C (encoding amino acids 541-666) and 66.1C (encoding amino acids 559-741). Both contained all or part of the coiled-coil region of Nischarin identified using the SMART motif prediction program (31, 32) (Figure 1A). The coiled-coil region corresponds to amino acids 624-694 in Nischarin and has the potential to engage in protein-protein interactions as well as to promote Nischarin homo-oligomerisation (33). To test whether the coiled coli region is necessary or sufficient for binding to ShcD, further Nischarin constructs were generated and tested for interaction with ShcD-CH2 domain using the yeast two hybrid assay.

**Figure 1.**
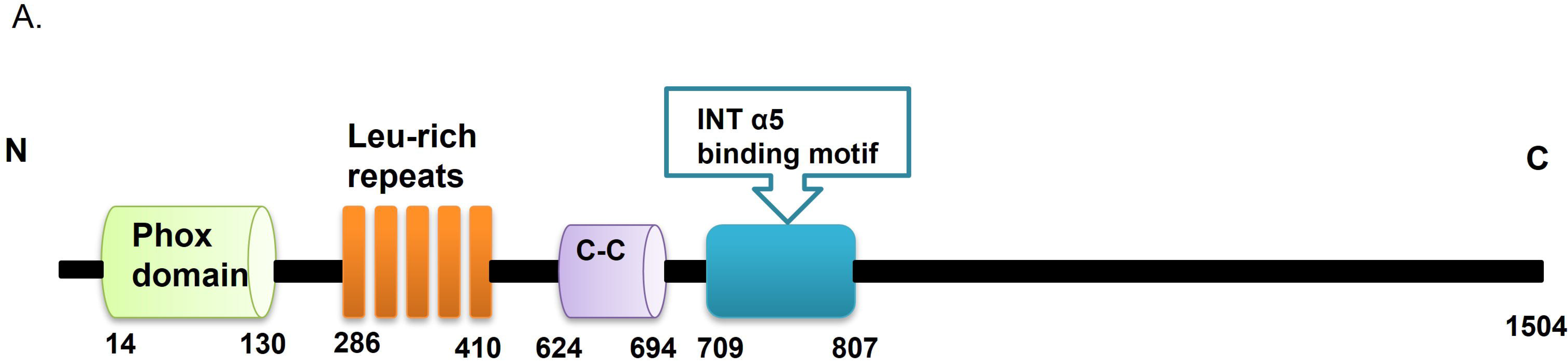
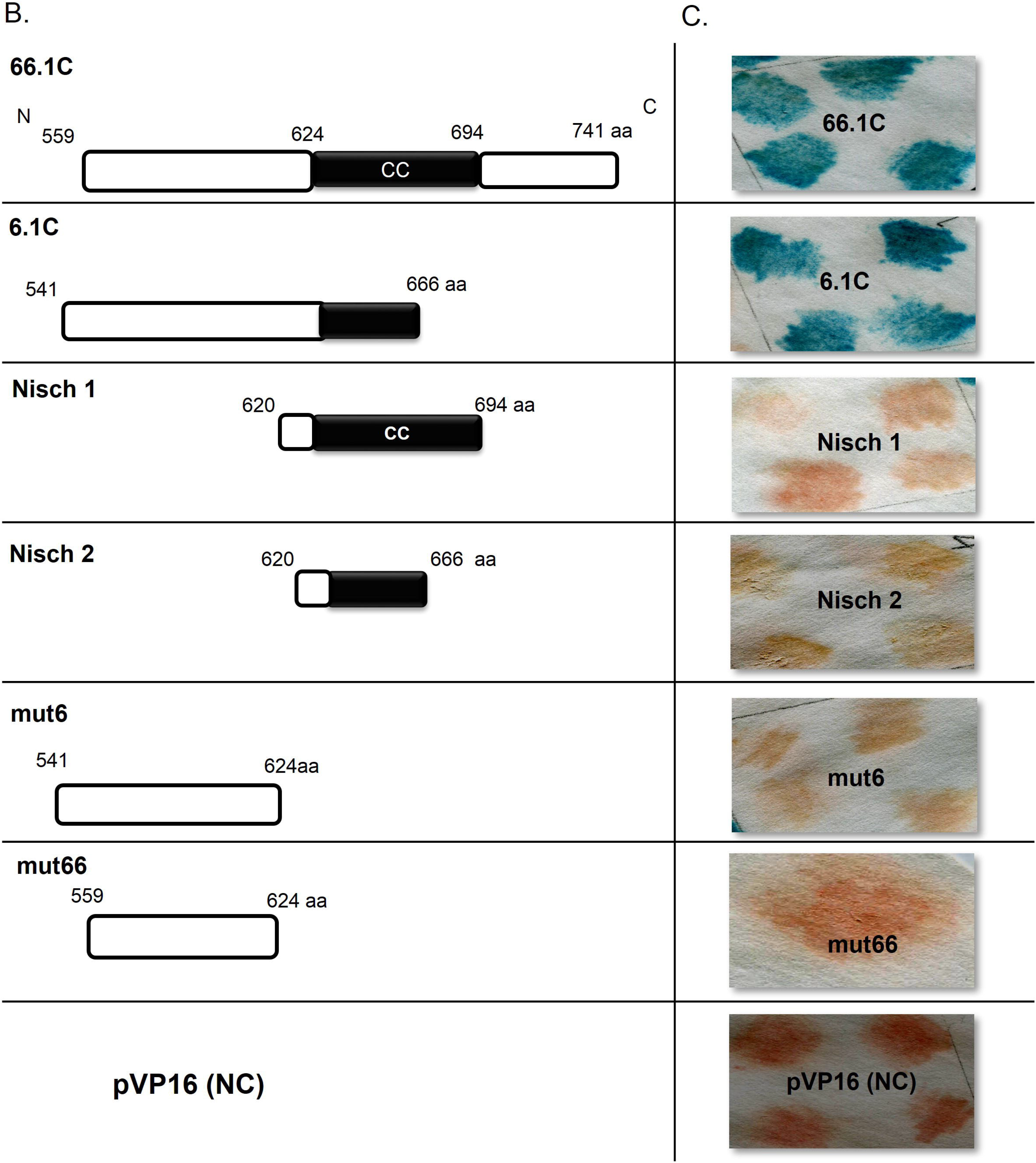
Yeast Two hybrid assays show the interaction between ShcD and Nischarin. (A) A schematic representation shows Nischarin structure that have been experimentally proved, adapted from (Sun Z. Et al, 2007; Swiss Prot 2008 accession number Q80TM9). C-C: Coiled-Coiled; N: amino terminus; C: carboxy terminus of the protein. (B) Nischarin sequences that were used in the two hybrid assays. (C)Yeast two hybrid β-galactosidase assay. After yeast transformation with prey and bait DNAs, yeast transformants were plated in a selective medium that lacked tryptophan and leucine. Thereafter, four colonies were picked and streaked on filter paper grown on top of a selection plate. After a couple of days, the cultured yeast was examined with the β-galactosidase assay for successful bait and prey interaction. The presence of the interaction is indicated by the blue colouration.

A stop codon was introduced into both clones 6.1C and 66.1C to produce the truncated proteins mut6 and mut66, which lacked the coiled-coil domains. Two additional constructs were generated; Nisch 1 contained the complete coiled-coil domain (amino acids 620-694), and Nisch 2, included the coiled-coil domain encoded by 6.1C (620–666) (Figure 1B).

β-Galactosidase transcription was detected by incubating yeast colonies with X-gal, which changes the yeast’s colour from pink to blue. The only colonies that turned blue were colonies that had been transformed with the original templates 6.1C and 66.1C together with ShcD-CH2 domain (Figure 1C).

Therefore, these data show that the coiled-coil region (amino acids 624-694) was not sufficient to mediate the ShcD/Nischarin interaction; however, this region was required, as mut 6 and mut66 did not interact with the ShcD-CH2 domain. Because both 6.1C and 66.1C interacted with ShcD-CH2, the interacting region was likely contained within the overlapping region from 559-666.

### Mapping the Nischarin interacting region on the CH2 domain of ShcD

To identify which region on the CH2 domain of ShcD is required to interact with Nischarin, FLAG-ShcD constructs with different truncations of the amino-terminal region, FLAG-ShcDΔ1-24 and FLAG-ShcDΔ1-93, were generated (Figure 2A). These correspond to the previously reported p69 and p59 versions of ShcD (3, 4). Each of the truncated versions and the full-length FLAG-ShcD was co-expressed with GFP-Nischarin in HEK 293 cells (Figure 2B). Anti-FLAG immunoprecipitates from the cell extracts were examined for the presence of Nischarin. Loss of the first 24 amino acids reduced the association of ShcD and Nischarin, while loss of the first 93 amino acids resulted in complete abrogation of ShcD and Nischarin interaction (Figure 2B). These data indicated that the Nischarin interaction region on ShcD is within the first 93 amino acids at the extreme amino terminus, and the first 24 amino acids contribute significantly to the interaction.

**Figure 2.**
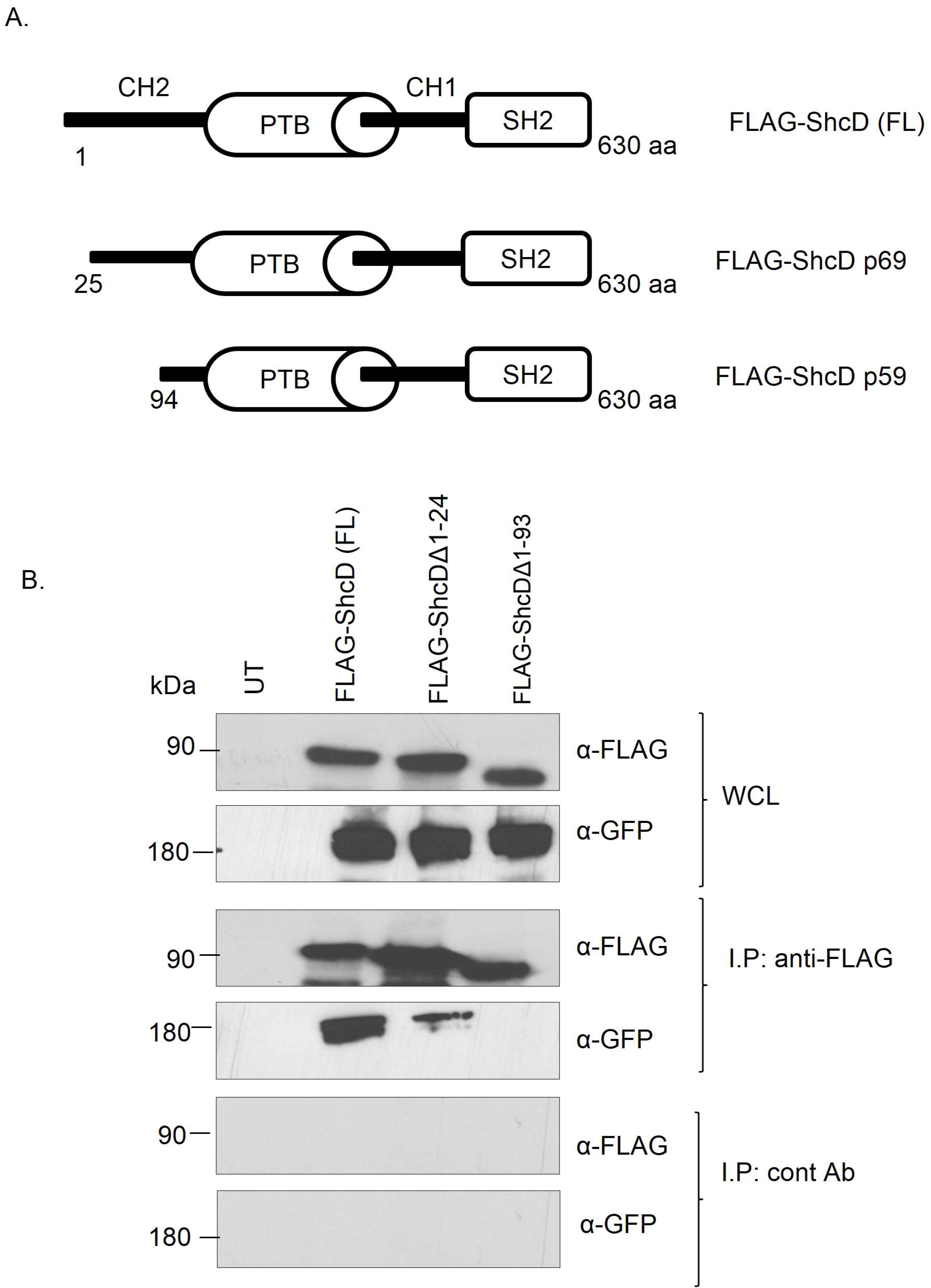
Mapping ShcD-CH2 domain for the Nischarin interacting region. (A) Schematic illustration demonstrates ShcD-CH2 truncated versions that were tested for their interaction with Niscahrin. (B) HEK 293 cells were transiently co-transfected with GFP-Nischarin and either FLAG-ShcD 1-630 (FL), FLAG-ShcDΔ1-24 (p69ShcD) or FLAG-ShcDΔ1-93 (p59 ShcD). The cells were lysed and incubated with 3 μg of immobilised anti-FLAG antibody or anti-PKTAG antibody (control) O/N at 4°C. The immunoprecipitated proteins were resolved on an 8% SDS-PAGE gel along with 1/10 of the whole cell lysate (WCL). Immunoblotting was performed with anti-FLAG and anti-GFP antibodies.

### ShcD co-immunoprecipitated with Nischarin in the MCF7 and MM253 cell lines

To study the functional impact of the Nischarin/ShcD interaction, we first decided to verify their interaction in both MCF7 and MM253 cell lines using co-immunoprecipitation. MCF7 and MM253 cell lines were either untransfected or transfected with different plasmid combinations: FLAG-ShcD+GFP, FLAG+GFP-Nischarin, or GFP-Nischarin+FLAG-ShcD. The expression of the recombinant proteins was tested by separating the whole cell lysates (WCLs) (Figure 3A and C), which was followed by co-immunoprecipitation in which beads were coupled to anti-FLAG antibody and incubated with the lysates obtained from MCF7 and MM253 cells subjected to different treatments. Finally, the immunoprecipitated proteins were immunoblotted with anti-GFP antibody and then re-probed with anti-FLAG. Western blot analysis suggests that FLAG-ShcD was able to pull down GFP-Nischarin in the MCF7 and MM253 cell lines, while the vector containing the FLAG tag alone was unable to immunoprecipitate Nischarin (Figure 3B & D). Therefore, FLAG-ShcD could immunoprecipitate GFP-Nischarin in MCF7 and MM253 cell lysates.

**Figure 3.**
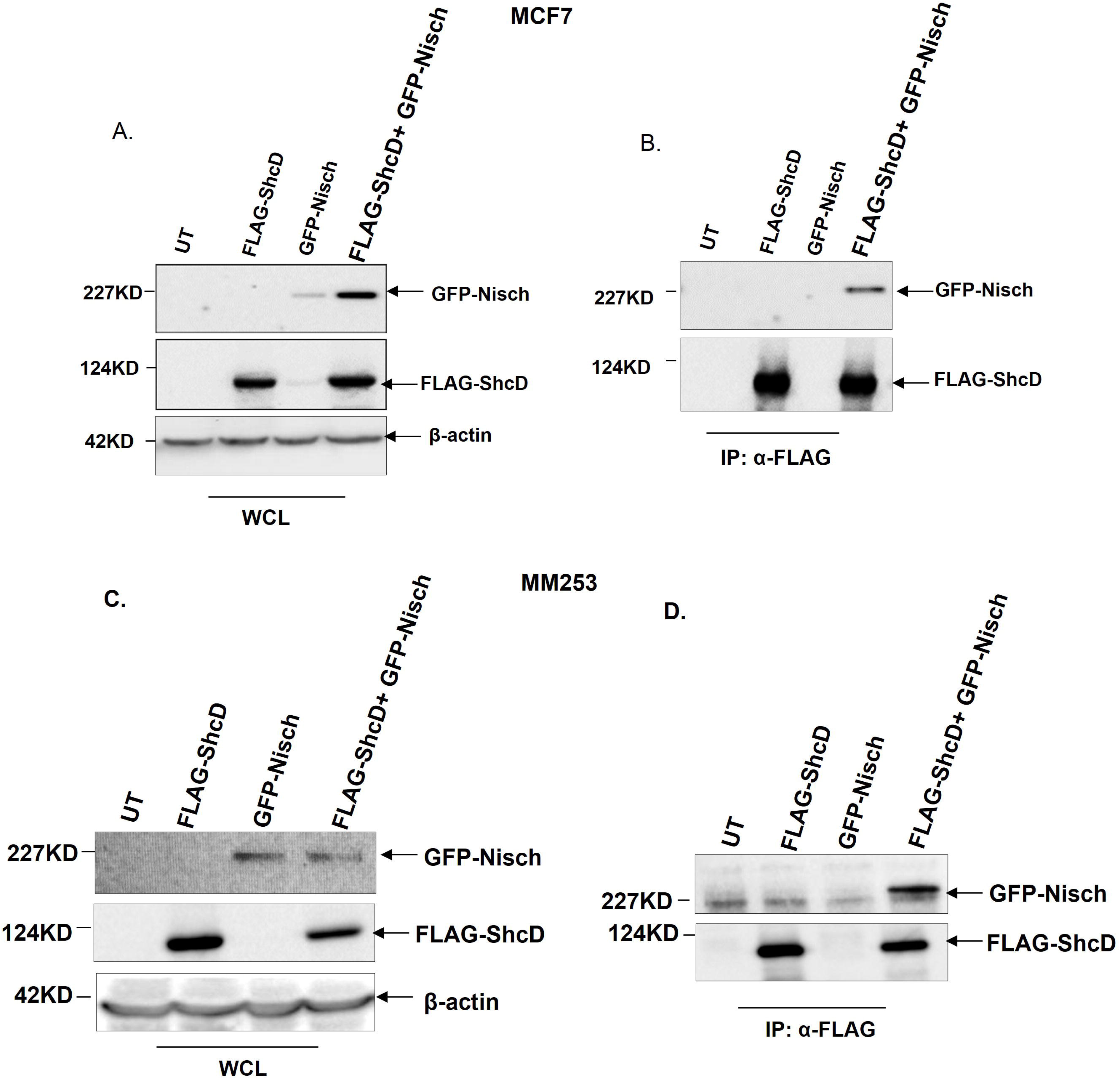
FLAG-ShcD co-immunoprecipitates with GFP-Nischarin from MCF7 and MM253 cell lysates. MCF7 and MM253 cells were seeded in 10 cm dishes. Next, the cells were either untransfected (UT) or transfected for 24 hrs with different plasmid combinations as follows: (GFP+FLAG-ShcD/FLAG-ShcD), (FLAG (-ve)+ GFP-Nischarin/GFP-Nisch), and (FLAG-ShcD+GFP-Nischarin/FLAG-ShcD+GFP-Nisch). Whole cell lysates (WCLs) from these cells were separated on an 8.5% SDS-PAGE gel. After proteins were transferred to the membrane, it was then immunoblotted with anti-GFP, anti-FLAG, and antibodies (A, C). The remaining cell lysates that were obtained from the untransfected and transfected cells were incubated with beads coupled to 2 µg of anti-FLAG antibody for 2 hrs. Next, the beads were washed, and the co-immunoprecipitated proteins were resolved on an 8.5% SDS-PAGE gel, transferred to PVDF membranes, and immunoblotted with anti-GFP and anti-FLAG antibodies (B, D). IP; Immunoprecipitation. UT; untransfected.

After proving that FLAG-ShcD immunoprecipitated with GFP-Nischarin in MCF7 and MM253 cell lines, we aimed to confirm that Nischarin also co-immunoprecipitates with ShcD by utilizing GFP-Trap, which comprises agarose beads that are covalently bound to anti-GFP antibody. MCF7 and MM253 cells were either untransfected or transfected with FLAG-ShcD+GFP, FLAG+GFP-Nischarin, or GFP-Nischarin+ FLAG-ShcD. Our data showed that GFP-Nischarin pulled down FLAG-ShcD in MCF7 and MM253 cells, whereas GFP was unable to co-immunoprecipitate FLAG-ShcD (Figure 4A and B). Thus, we deduced that Nischarin can co-immunoprecipitate with ShcD in the lysates obtained from the co-transfected MCF7 and MM253 cells. While the yeast two hybrid data suggest that the interaction between the ShcD-CH2 domain and Nischarin is direct, it is quite possible that in the immunoprecipitations, other proteins may be involved in stabilising the complex.

**Figure 4.**
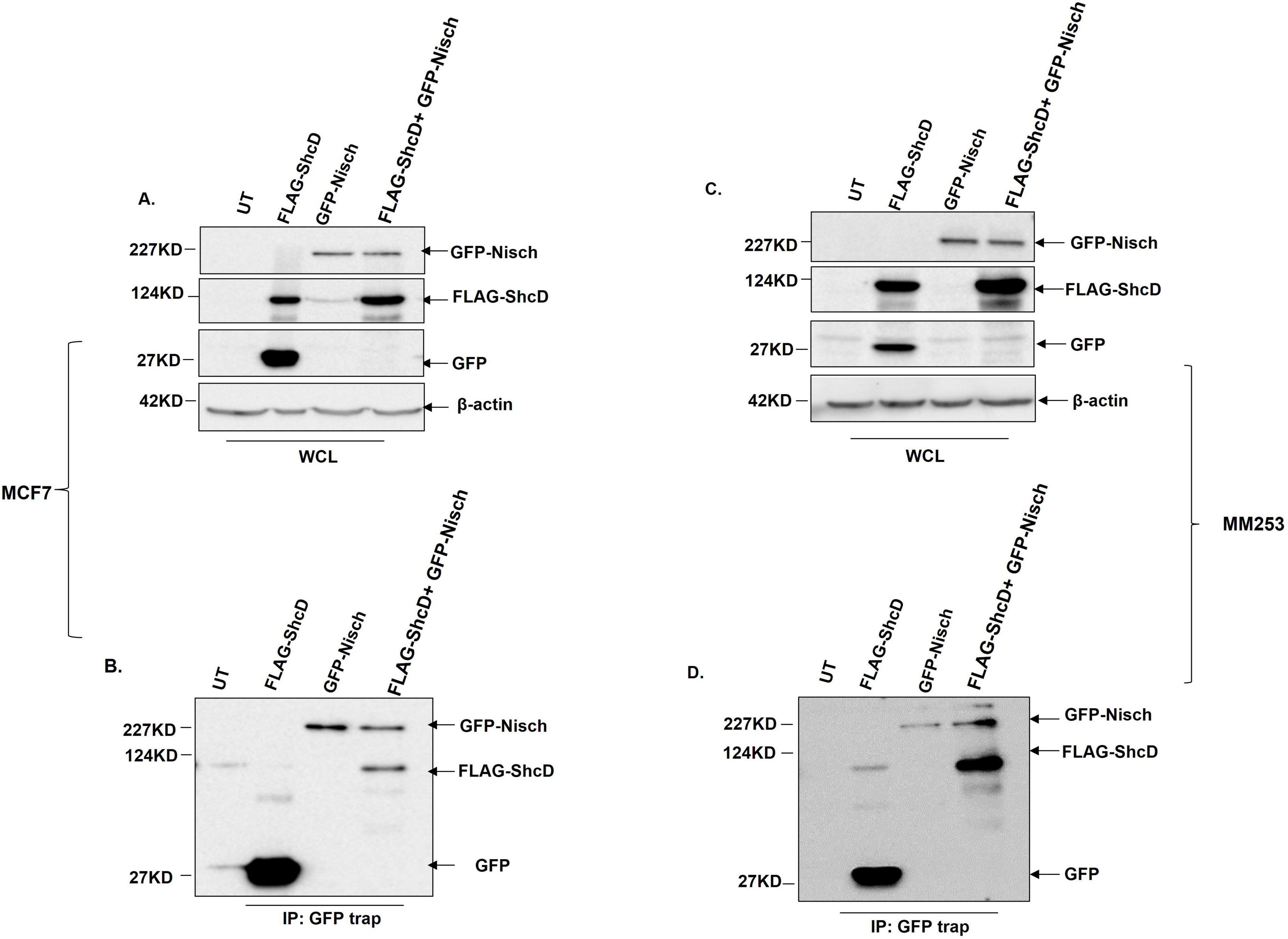
GFP-Nischarin co-immunoprecipitated with FLAG-ShcD from MCF7 and MM253 cell lysates. MCF7 and MM253 cells were seeded in 10 cm dishes. After 24 hrs, the cells were either untransfected (UT) or transfected for 24 hrs with different plasmid combinations, as follows: (GFP+FLAG-ShcD/FLAG-ShcD), (FLAG (-ve) +GFP-Nischarin/GFP-Nisch), and (FLAG-ShcD+GFP-Nischarin/FLAG-ShcD+GFP-Nisch). The cells were then lysed. The whole cell lysates (WCL), A and C, were resolved on 8.5% SDS-PAGE gel and immunoblotting was performed using anti-GFP, anti-FLAG and anti-β-actin antibodies. The rest of the lysates were incubated with GFP trap overnight at 4□C. Next day, the beads were washed and the immunoprecipitated proteins were resolved on SDS-PAGE gel. Immunoblotting was then performed and the with anti-GFP and anti-FLAG antibodies, B and D.

### Confirming the cellular colocalisation of ShcD and Nischarin by immunofluorescence microscopy

To further investigate the interaction between Nischarin and ShcD, we decided to study their subcellular localisation using fluorescence microscopy. MCF7 and MM253 cells were transfected with GFP-Nischarin + FLAG, GFP + FLAG-ShcD, GFP + FLAG, or GFP-Nischarin + FLAG-ShcD. Both Nischarin and ShcD exhibited predominant cytoplasmic distribution (Figure 5A and B). Co-localisation was evaluated by merging the resulting images using ImageJ-win64 software, which revealed a clear overlap between Nischarin (green channel) and ShcD (red channel) in both MCF7 and MM253 cells. Their co-localisation was observed in the cytoplasm as well as at a membrane-like structure, particularly in MM253 cells (Figure 5B). Nischarin and ShcD co-localisation provides more evidence for the interaction between the two proteins.

**Figure 5.**
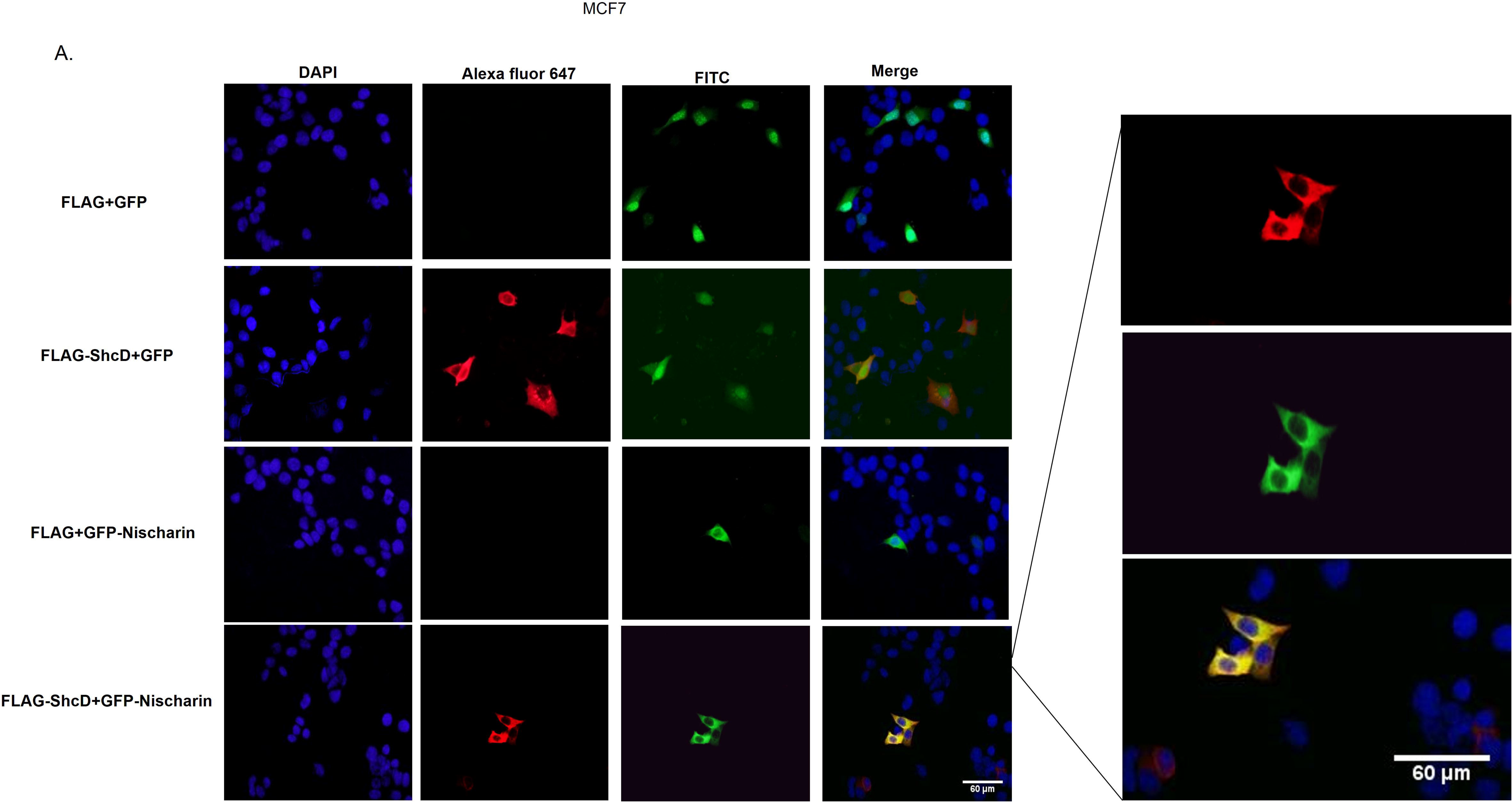
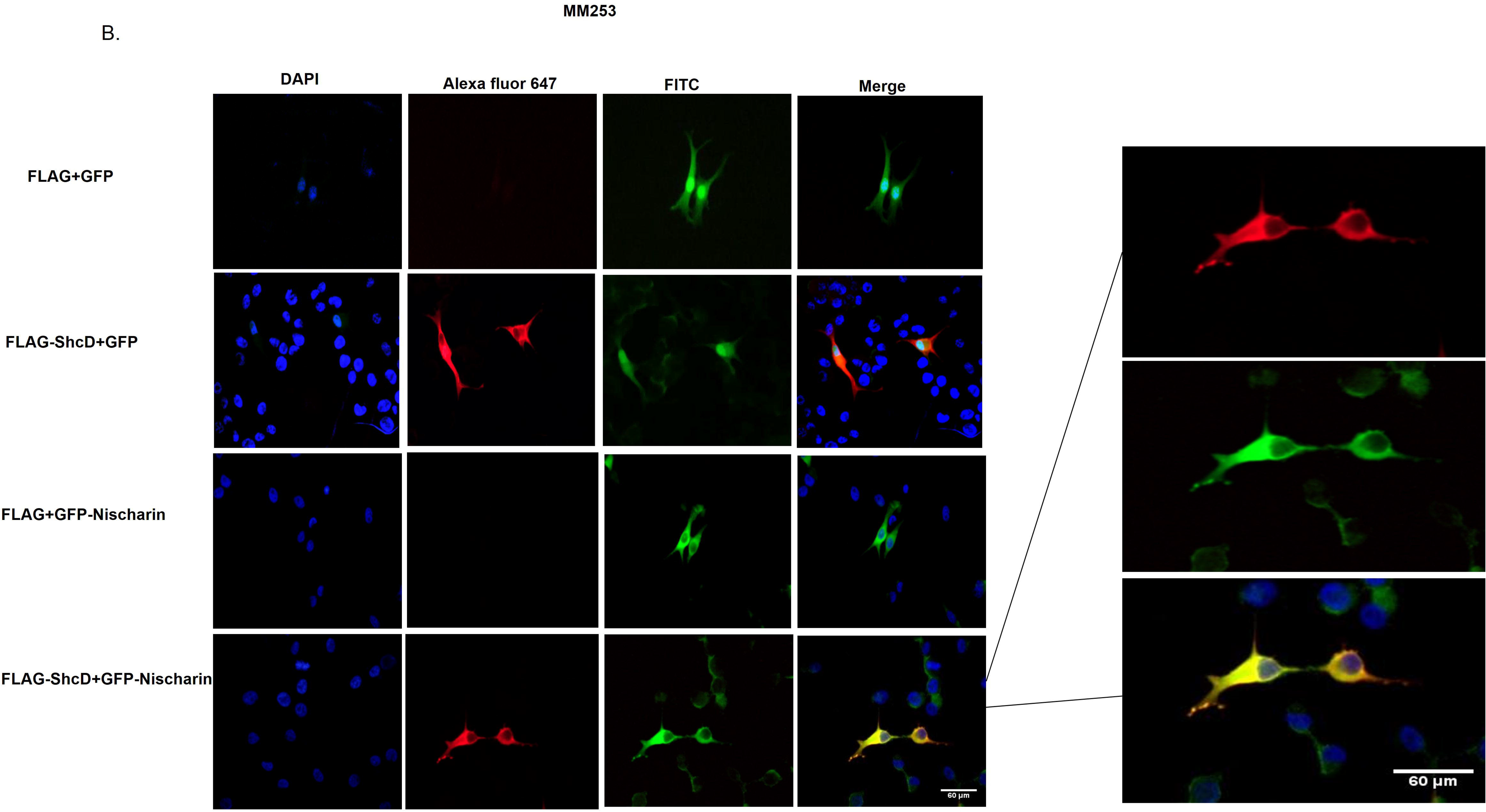
Subcellular co-localisation of ShcD and Nischarin in MCF7 and MM253 cell lines. MCF7 (A) and MM253 (B) cell lines were seeded on cover slips in 6-well plates. Next, cells were transfected with the following combinations of plasmids: (GFP-Nischarin+ FLAG), (GFP+ FLAG-ShcD), (GFP+FLAG), and (GFP-Nischarin +FLAG-ShcD). After 24 hrs of transfection, cells were fixed, permeabilised, blocked, and incubated with primary antibody (anti-FLAG). Immunofluorescence was visualized on a fluorescence microscope (Olympus, BX51TF) and analysed with cellSens Standard software. A 40x objective was used. Co-localisation was evaluated by merging the resulting images using ImageJ-win64 software. Zoomed images represent the overlap of the green and red channels at the membrane area. Scale bar: 60 µm.

### The co-expression of Nischarin and ShcD attenuates ShcD-mediated LIMK phosphorylation

Since Nischarin was previously shown to inhibit the phosphorylation of LIM kinase (LIMK), thereby inhibiting cell motility, we aimed to investigate the impact of Nischarin and ShcD co-expression on phospho-LIMK levels. We used recombinant LIMK because the anti-phospho-LIMK antibody available to us only recognises recombinant LIMK. MCF7 and MM253 cells were transfected with different combinations of plasmids, and the resulting cell lysates were immunoblotted to explore phospho-LIMK levels in the different transfection conditions (Figure 6A and B). As expected, phosphorylated LIMK levels were remarkably decreased in both cell lines overexpressing Nischarin, while ShcD overexpression alone resulted in a slight increase in phosphorylated LIMK, which was more noticeable in MM253 cells than in MCF7 cells (Figure 6A and B). Unexpectedly, cells co-expressing both Nischarin and ShcD demonstrated reduced levels of phospho-LIMK. Thus, when ShcD and Nischarin were co-expressed in MCF7 and MM253 cell lines, the effect of Nischarin on phospho-LIMK dominated that of ShcD.

**Figure 6.**
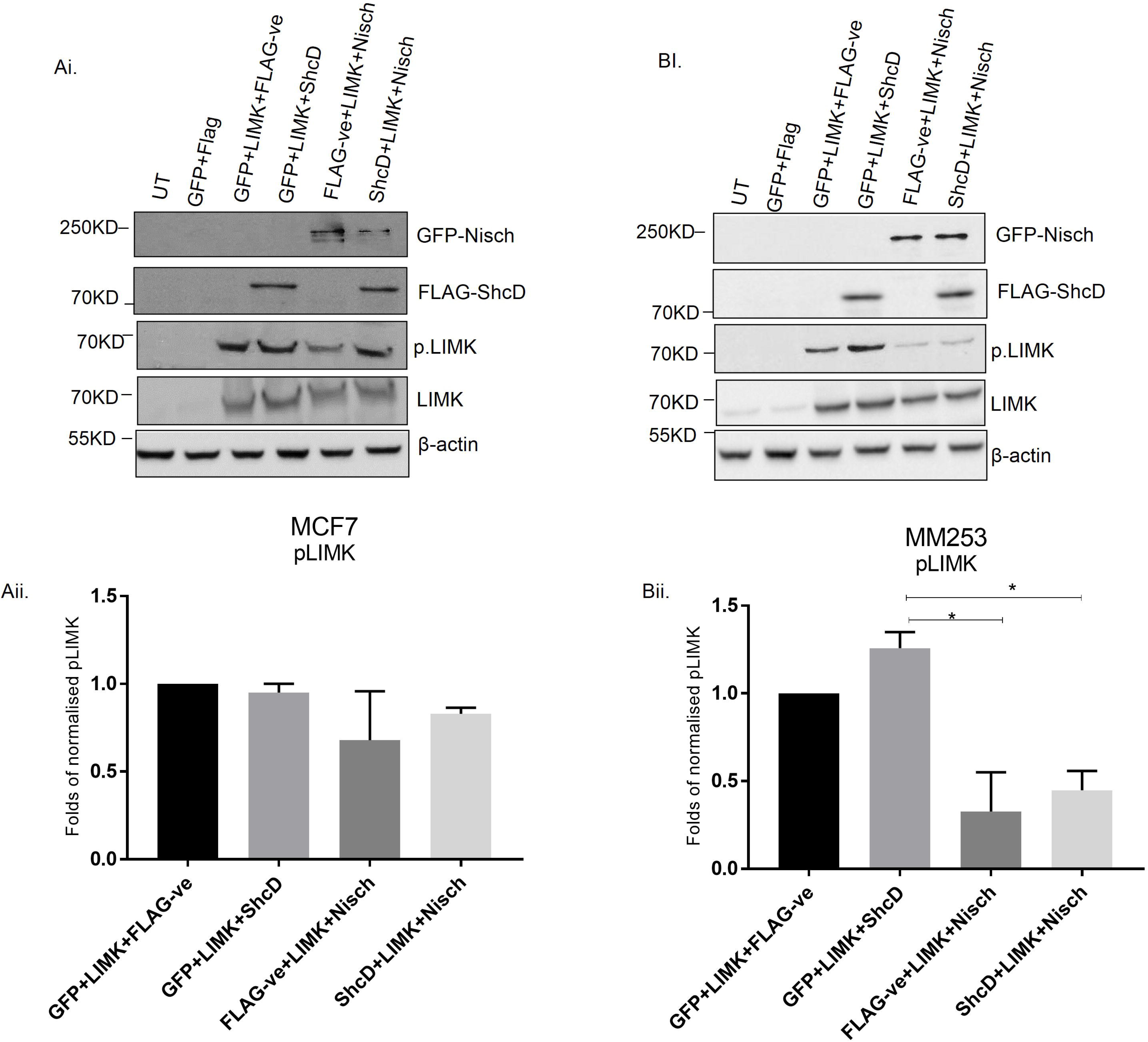
The impact of the association of the ShcD adaptor protein and Nischarin cytosolic protein on the phosphorylation of LIMK. MCF7 cells (Ai) and MM253 (Bi) cells were seeded in 6-well plates. The next day, cells were untransfected (UT) or transfected with LIM kinase for 24 hrs. The cells were then lysed, and proteins were separated by SDS-PAGE though an 8.5% gel and transferred onto PVDF membranes. The membranes were first probed with anti-LIMK antibody and then re-probed with β-actin antibody. Aii and Bii are pool of three independent experiments for MCF7 and MM253, respectively. Error bars are represented by SEM. Bands intensities were analysed using ImageJ.

### The co-expression of Nischarin and ShcD weakens ShcD-mediated ERK phosphorylation

Nischarin was reported to negatively regulate phosphorylated ERK levels (34). Therefore, we aimed to determine the effect of Nischarin and ShcD on phosphorylated ERK. MCF7 and MM253 cells were transfected as previously described in the LIMK experiment. Following 20 hrs of serum starvation, cells were cultured with complete medium for 30 min and levels of phosphorylated ERK were examined by western blotting. According to the normalized phospho-ERK band density analysis, ShcD-overexpressing cells demonstrated elevated levels of phospho-ERK in both cell lines, while Nischarin-overexpressing cells showed reduced levels of phospho-ERK. Similar to the LIMK experiment, ShcD failed to rescue the reduction in the levels of phospho-ERK mediated by Nischarin (Figure 7A and B). These results suggest that ShcD was unable to surmount the inhibitory effect of Nischarin on phospho-ERK. An unexpected insignificant elevation of phospho-ERK was observed in GFP transfected MCF7 cells.

**Figure 7.**
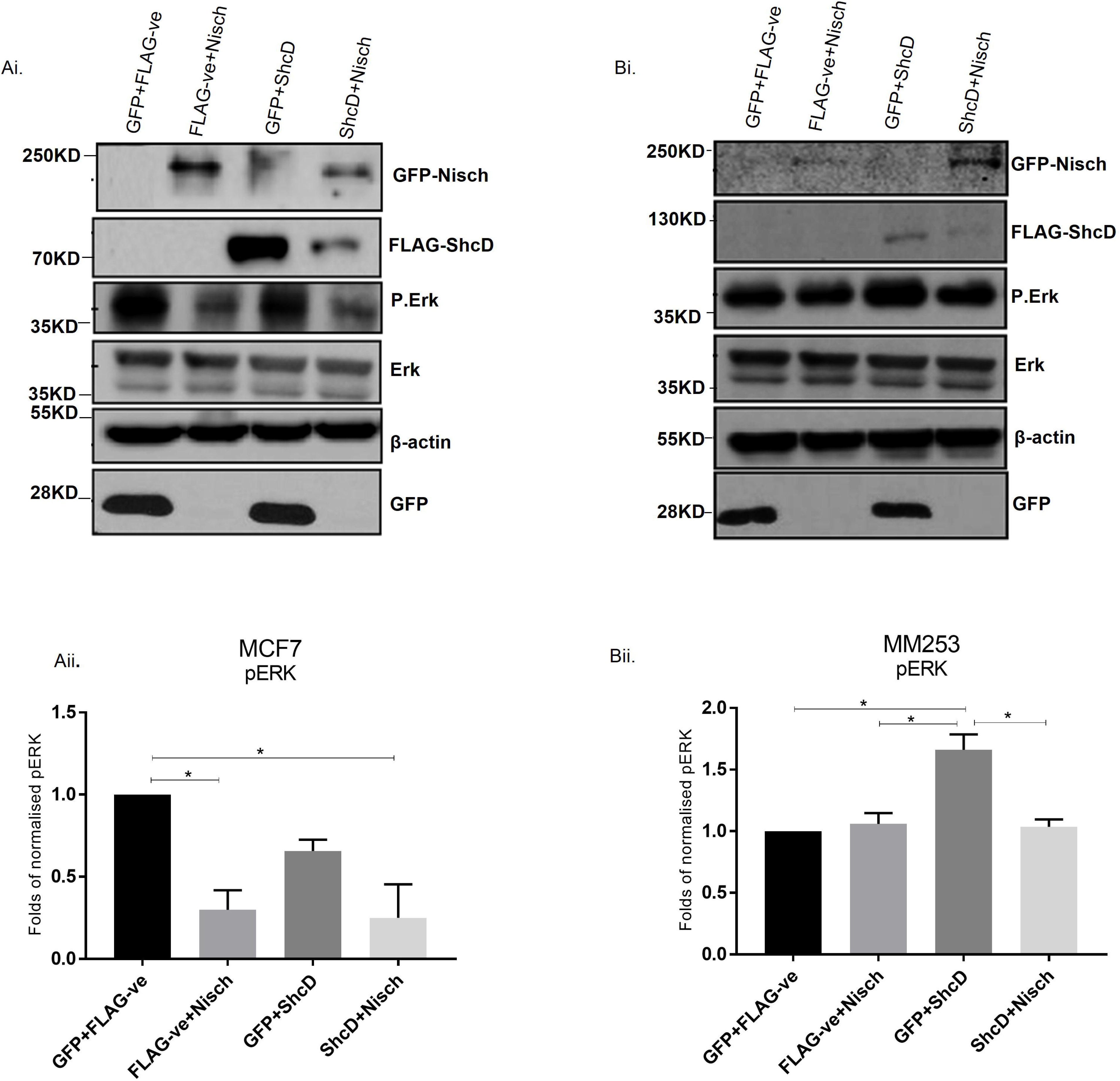
The impact of the association of the ShcD adaptor protein and Nischarin cytosolic protein on the phosphorylation of ERK. MCF7 cells (Ai) and MM253 cells (Bi) were seeded in 6-well plates. Next, they were transfected with GFP+FLAG, GFP+FLAG-ShcD, FLAG+GFP-Nischarin, or GFP-Nischarin+ShcD. After 24 hrs of transfection, cells were starved using RPMI starvation media (0.1% FBS). Following 24 hrs of starvation, cells were treated with complete media for 30 mins and lysed. Then, proteins were resolved on an 8.5% SDS-PAGE gel and transferred to a PVDF membrane. Next, the membrane was immunoblotted with anti-phospho-ERK and re-probed with anti-ERK, anti-FLAG, anti-GFP, and finally β-actin. Aii and Bii are pool of three independent experiments for MCF7 and MM253, respectively. Error bars are represented by SEM. Bands intensities were analysed using ImageJ.

### Nischarin negatively impacts ShcD-induced cell motility

After investigating the effect of Nischarin and ShcD on the phosphorylation of ERK and LIM kinase, it was vital to test the functional impact of this interaction on the migration of MCF7 and MM253 cells. For this purpose, MM253 and MCF7 cells were transfected as described above. Next, transwell and wound-healing migration assays were performed. MM253 or MCF7 cells were allowed to migrate with a serum gradient from the upper chamber to the lower chamber through a porous membrane for 24 hrs. Cells that migrated to the membrane face in the lower chamber were stained with crystal violet and then lysed with 4% SDS; the absorbance readings at 570 nm were acquired on a microplate reader. The percentage of migrated cells was calculated and is presented in bar charts. The concomitant expression of Nischarin and ShcD significantly reduced the latter’s ability to enhance cell migration (p<0.05) (Figure 8A and B). To further validate the results obtained from the transwell assays, wound-healing assays were conducted. Twenty-four hours after transfection, a wound was made across the cell monolayer, and images were taken at 0 hrs and 24 hrs after scratching. MCF7 cells transfected with ShcD showed an increased rate of migration, whereas cells with Nischarin and ShcD co-expression exhibited less migratory ability, as indicated by the extent of wound closure (Figure 8C).

**Figure 8.**
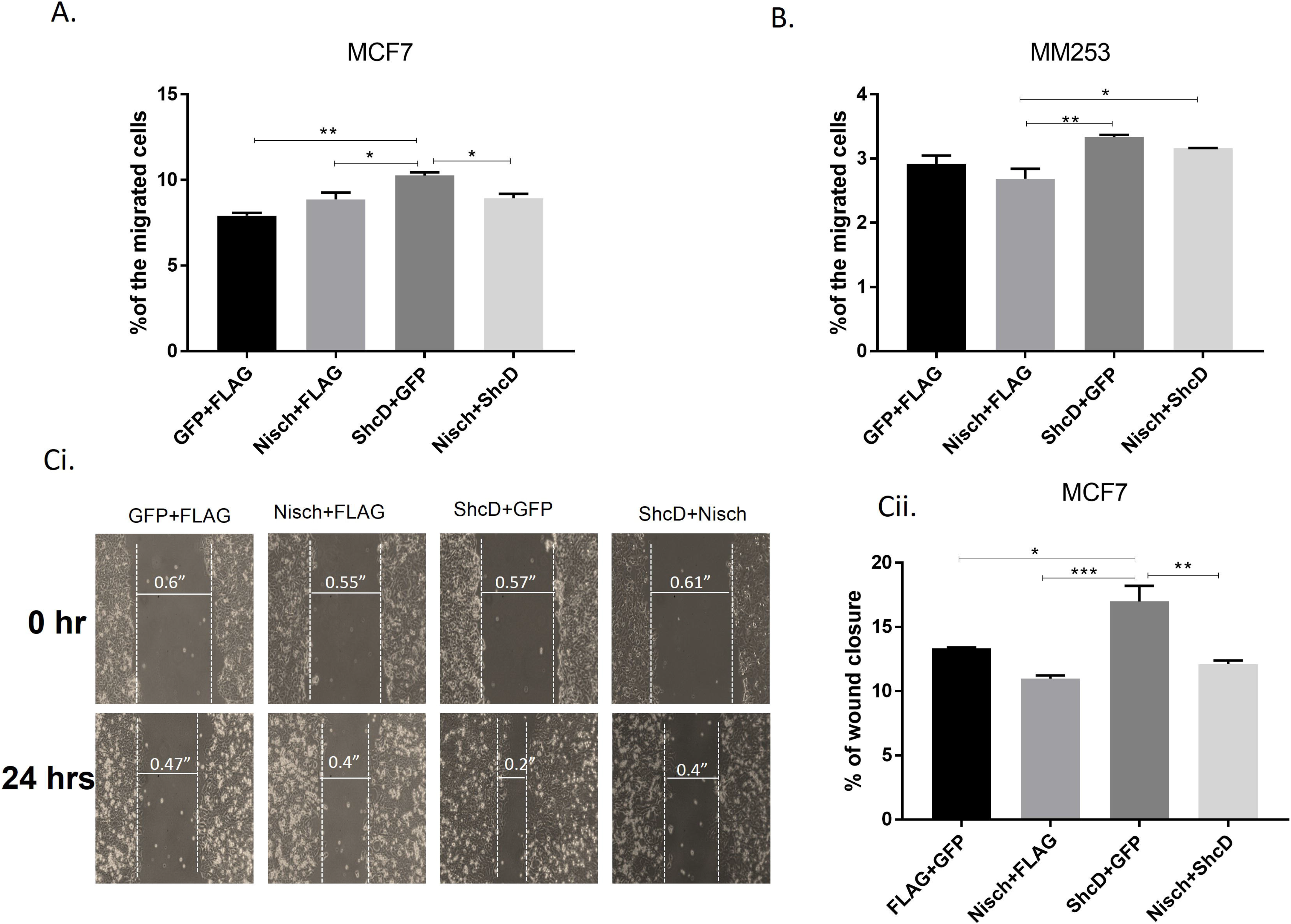
The effect of the association of ShcD and Nischarin on cell migration. (A, B) MCF7 and MM253 cells were transfected with the following combinations of DNA plasmids: (GFP+FLAG), (GFP-Nischarin+FLAG), (GFP+FLAG-ShcD), and (GFP-Nischarin+FLAG-ShcD). Cells (1.25x10^5^) from each transfection set were resuspended in 0.1% serum and added to the upper chamber of a Boyden chamber. The cells were allowed to migrate across the porous membrane towards media supplemented with 10% serum for 24 hrs. The migrated cells were stained with crystal violet, and the colourimetric readings were taken at 570 nm. Scale bars represent SEM. (Ci) MCF7 cells were transfected as described in A and B. A scratch was made through the cell monolayer, and images of the wounds were taken at 0 and 24 hrs after scratching using an Optika microscope with a 10X objective. (Cii) The bar chart represents a pool of 2 independent wound healing experiments in which each condition was imaged from three different fields.

### Nischarin expression is associated with increased overall survival of patients with metastatic breast cancer based on publicly available data

Since the Nischarin and ShcD association had a negative impact on migration, it was of interest to explore the impact of this association on the OS of breast cancer patients. To this end, patient data relating to Nischarin and ShcD expression patterns were collected utilizing bioinformatics databases bc-GenExMiner 4.2 database tool and KM plotter database (27–29). This was followed by dividing the patients into two groups based on the lymph node (LN) status: lymph node positive (LN +ve) and lymph node negative (LN-ve). Kaplan-Meier plot curves for survival were plotted. According to the OS curves, metastatic breast cancer patients (LN-ve) with ShcD overexpression displayed a decreased survival rate, whereas opposing findings were observed in patients with Nischarin expression (Figure 9A). Patients with high expression of both proteins had better OS (*p=0.029<0.05*) than did patients with high ShcD expression alone (*p=0.047<0.05*). The same effect was observed in (LN+ve) breast cancer patients with individual expression of Nischarin and ShcD; nevertheless, co-expression of both proteins corresponded to better OS (*p=0.17>0.05*). The OS and RFS of all patients (LN-ve and LN+ve) who had high expression of both Nischarin and ShcD demonstrated increased survival rates (*p=0.0017<0.01 and p<1E-16<0.01*, respectively) compared to those with high expression of ShcD alone (Figure 9B).

**Figure 9.**
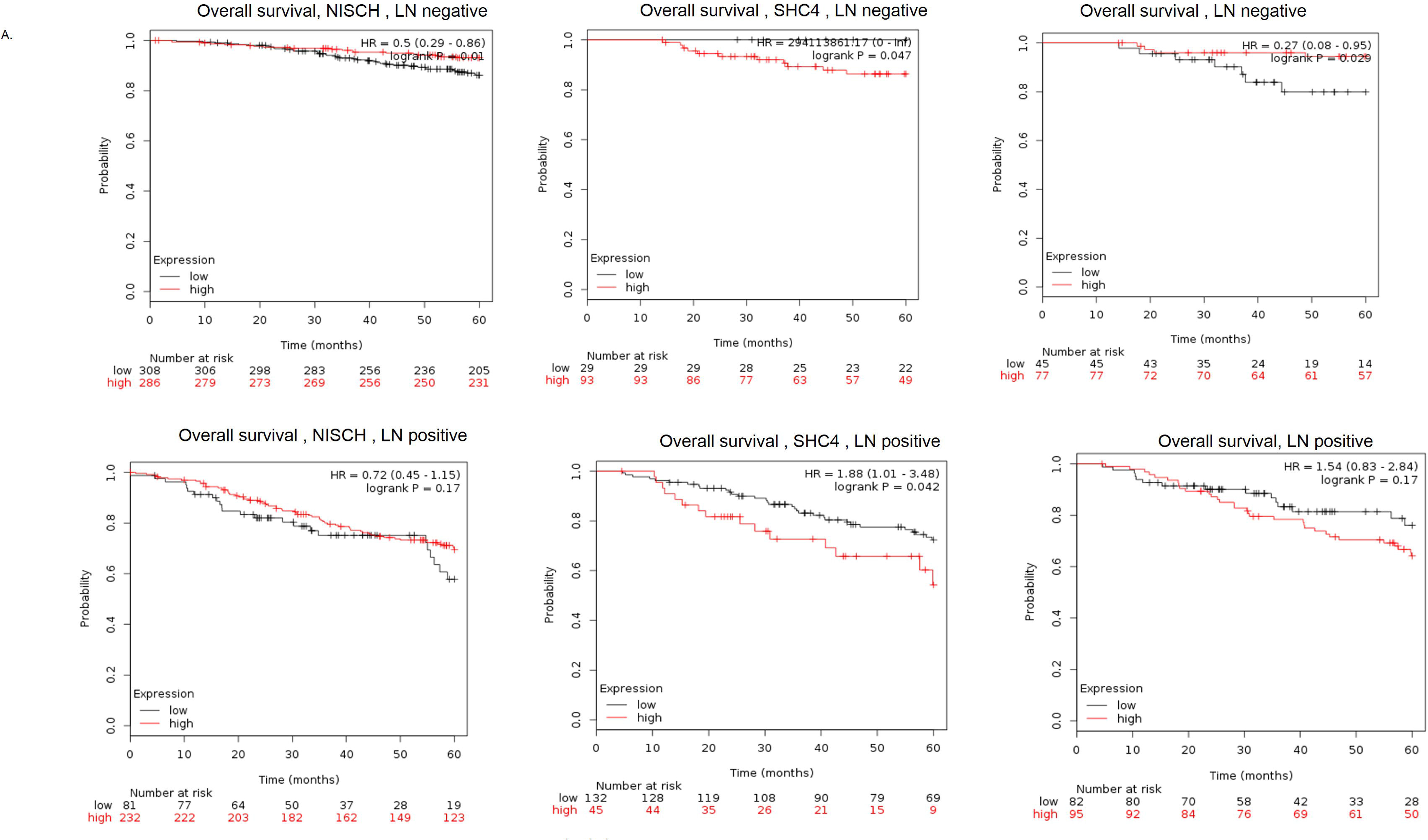
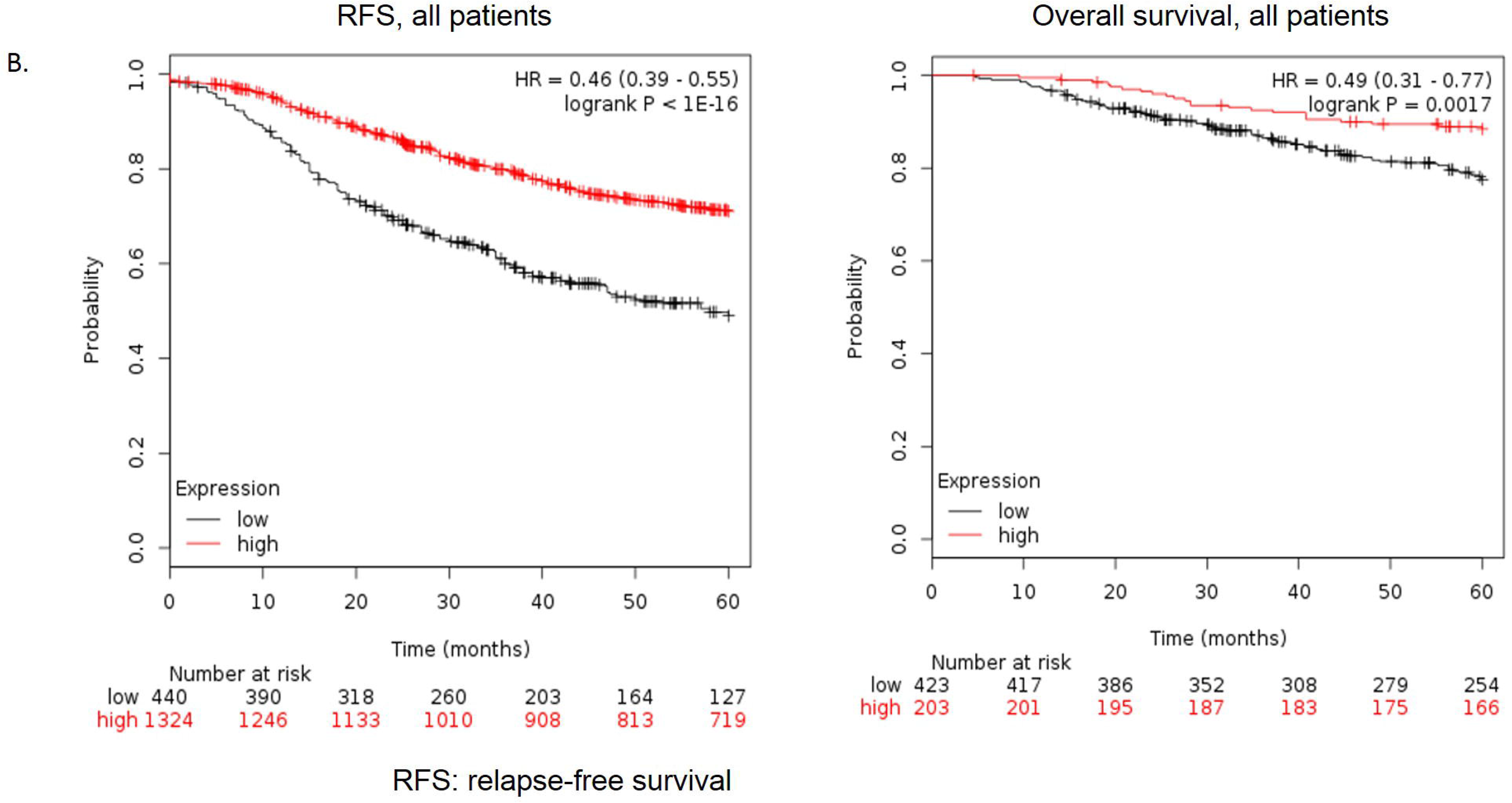
In silico analysis of data from breast cancer patients with differential expression of ShcD and Nischarin. (A) Kaplan-Meier survival curves for breast cancer patients with overexpression of ShcD/4, Nischarin (Nisch), or both according to lymph node (LN) status (i.e., LN-ve or LN+ve). (B) Survival curves illustrate the effect of Nischarin and Shc4 concomitant expression on the overall survival (OS) and relapse-free survival (RFS).

## Discussion

In 2007, ShcD was first identified as a positive regulator to melanoma cell migration as well as it was found to be upregulated in 50% of invasive and metastatic melanomas (3). Since then, few studies experimented the role of ShcD in cell migration (12, 35, 36). Therefore, in this study, it was aimed to further investigate ShcD role in cellular motility. The followed approach was to identify potential interacting proteins that may contribute to ShcD migratory role; therefore, a yeast two hybrid screen was conducted. Interestingly, the invasion suppressor protein, Nischarin was identified as a binding partner for the ShcD-CH2 domain.

The collagen homology domains (CH1/2) are the least conserved among Shc proteins (2); hence, it was of interest to explore whether the ShcD-CH2 domain exhibits a unique interaction with Nischarin and not the CH2 domains possessed by the other Shc members. Based on the results of the UniProt alignment program, the percentage amino acid sequence identity between the ShcD-CH2 domain and p66ShcA-CH2 domain is 23%, while it is 21% for ShcC and 14% for ShcB. Our data showed that Nischarin associates with ShcD but not with p66ShcA, a member from the same family that contains a CH2 domain (Figure 2S).

Prior to investigating Nischarin and ShcD interaction in mammalian cell lines, it was decided to identify the Nischarin interacting sequence that is required for ShcD binding. Nischarin is a large cytosolic protein comprising several domains that enable it to interact with different proteins (33). The coiled-coil domain in Nischarin represents one of the domains that facilitate protein-protein interactions (33). Therefore, it was speculated that this Nischarin coiled-coil region might be sufficient to mediate the Nischarin and ShcD interaction. Utilising yeast two hybrid assay technique, Nischarin coiled-coil region was found to be insufficient to mediate its interaction with ShcD. One study indicated that Nischarin or IRAS, the human homologue of Nischarin, can homo-oligomerise through the coiled-coil region (33). Presumably, the coiled-coil motif is likely to be required for Nischarin homo-oligomerisation, leading to the formation of Nischarin homodimers. Two copies of the interacting region of Nischarin may be required to be in close proximity to facilitate ShcD interaction, a phenomenon that requires further experimentation.

It was important to examine this interaction in mammalian cells to validate the obtained results from the yeast two hybrid assays as well as to elucidate the importance of the ShcD-CH2 in this interaction. Co-immunoprecipitation experiments using full length and truncated versions of ShcD provided clear evidence that full length ShcD and Nischarin interacted in mammalian cells. ShcD lacking the amino-terminal CH2 domain failed to immunoprecipitate Nischarin; therefore, the ShcD-CH2 domain is required for interaction with Nischarin. The CH2 domain was further mapped to identify the minimum region required for full binding. The region comprising amino acids 1 to 93 was indispensable for this interaction to occur; notably, deleting this sequence results in ShcD nuclear translocation (37). This finding might indicate that the cytoplasmic presence of ShcD is crucial for Nischarin binding. Deletion of amino acids 1 to 24 reduced the interaction, which suggests the importance of these residues for stabilising the interaction. For the first time, it was shown that the CH2 domain of ShcD is involved in protein-protein interactions.

Nischarin interacts with other proteins that are involved in cell motility, including α5β1 integrin, PAK, LIMK, Rac (19, 38, 39). Importantly, the Nischarin region involved in the ShcD interaction (amino acids 541-666) might overlap with the regions required for binding with PAK, LIMK and α5β1 integrin. It will be of interest to determine which of these proteins can compete for ShcD binding.

Nischarin functions as a tumour suppressor in breast cancer and as a negative regulator to cell motility (18–20, 34, 38, 40). To this end it was crucial to study the functional consequence of Nischarin-ShcD interaction. Prior to studying the functional consequence of Nischarin and ShcD in a breast cancer cell line (MCF7) and a melanoma cell line (MM253), it was crucial to prove their interaction in these two cell lines. Our results provided clear evidence that ShcD interacts with Nischarin in both cell lines.

The association was further evaluated by observing their co-localisation using fluorescence microscopy. ShcD and Nischarin were predominantly distributed in the cytoplasm, which is consistent with previous studies on both ShcD and Nischarin. Nischarin and ShcD co-expression showed clear evidence of their co-localisation in the cytoplasm as well as at the membrane region (3, 19).

It worth to mention that mouse Nischarin was used for the experiments in this study. Mouse Nischarin (mNischarin) Shows 80% homology with human Nischarin (hNischarin). The main difference is that mNischarin lacks the amino terminal domain, despite this difference unpublished data demonstrated that hNischarin does interact with ShcD, which indicates that the amnio terminal region in hNischarin is not required for ShcD and Niscahrin association.

To this end, it was of interest to explore the functional consequence of ShcD and Nischarin. Since Nischarin was reported to inhibit LIMK and ERK activation (34), we were prompted to examine the functional consequences of the Nischarin and ShcD interaction on the LIMK and ERK pathways. It was anticipated that ShcD might relieve the Nischarin inhibitory effect on LIMK and ERK, allowing the cells to acquire a more motile phenotype.

Consistent with previous studies, our findings showed that Nischarin overexpression inhibited both phospho-LIMK and phospho-ERK, while upregulation of ShcD promoted the phosphorylation of ERK and LIMK. Unexpectedly, ShcD failed to rescue the reduction in the levels of phospho-ERK and phospho-LIMK when it was co-expressed with Nischarin. Nischarin is known to inhibit different pathways by interacting with different proteins, such as LIMK, PAK, α5β1 integrin, ERK and FAK (18, 19, 34, 40). Nischarin inhibited the ability of ShcD to mediate ERK phosphorylation in MM253 cells but not in MCF7 cells. The failure of Nischarin to elicit the same effect in MCF7 cells could be because ShcD has a predominant effect on LIMK activation in these cells. In unpublished data, ShcD was observed to interact with LIMK regardless of the expression of Nischarin. Furthermore, in v-ErbB-transformed fibroblasts, ShcA was found to form a multi-phospho-protein complex containing PAK and MLCK, which are involved in LIMK signalling (41). The predominant effect of ShcD on phospho-LIMK could also be explained by the fact that ShcD might be involved in transducing different active signal pathways in MCF7 cells, such as those mediated by EGFR, IGF-1, and VEGFR, which have been proposed to interact with ShcD (3, 13, 42). A point worth to be mentioned is that GFP transfection in control MCF7 cells was found to result in ERK phosphorylation unlike MM253 melanoma cells. GFP transfection was proved to promote oxidative stress which might explain the slight elevation of phospho-ERK (43, 44). In contrary, MM253 cells did not show the same observation this could be explained by the fact that melanocytes are equipped with active antioxidant defences mechanisms (45).

Previous reports proved the involvement of ERK and LIMK in cell migration (46, 47). Since Nischarin and ShcD co-expression negatively affected ERK and LIMK phosphorylation, we were interested to determining whether this has further implications for cell motility. Nischarin-overexpressing cells exhibited relatively less migration than did ShcD-overexpressing cells, which is consistent with the role of Nischarin in cell migration. In contrast to cells overexpressing Nischarin, ShcD-overexpressing cells demonstrated higher migratory ability, which agrees with the reported role of ShcD in cell migration. Transwell and wound-healing migration assays showed that ShcD-mediated cell migration was suppressed when Nischarin was co-transfected with ShcD. This result describes the impact of ERK and LIMK inhibition on cell motility when Nischarin and ShcD are overexpressed.

In silico analysis of publicly available data of breast cancer patients revealed that patients with Nischarin expression have a better prognosis than do patients expressing ShcD. Patients with both Nischarin and ShcD expression showed better survival, which indicated that the tumour suppressor abilities of Nischarin could overcome the oncogenic activity of ShcD. Interestingly, lymph node positive patients with concomitant expression of Nischarin and ShcD showed poorer prognosis, although the p-value was not significant. Based on this observation, we assumed that the ratio between the expression of the two proteins determines the ultimate biological response; this observation needs further investigation.

Herein for the first time, we were able to show that Nischarin is an interacting partner of ShcD and could mitigate the effect of ShcD on cell migration by blocking ERK and LIMK phosphorylation. Although the mechanism by which Nischarin inhibits the effects of ShcD has yet to be determined, the functional impact of the Nischarin and ShcD interaction provides new insight into the role of ShcD in migration.

## Conclusion

A unique interacting partner of ShcD, Nischarin, was found to interact with ShcD-CH2 domain. The association of Nischarin with ShcD was found to negatively influence ShcD-mediated LIMK and ERK-phosphorylation as well as cell migration. Detailed molecular studies are required to gain further insight into the mechanism of how Nischarin adversely affects ShcD migratory function. The CH2 domain is likely involved in additional signals that should be examined to further elucidate the functions of the scarcely studied ShcD protein.

DTT 1,4-Dithiothreitol

## Limitations

The experiments were performed using transient transfection which does not ensure the uniform expression of the ShcD and Nischarin in all the experiments. One additional limitation, from our experience there is no reliable antibody for ShcD; therefore, we tend to use recombinant ShcD tagged to FLAG. It is worth to note that the role of ShcD in melanoma migration results from its overexpression, which supports the used system. The wound migration assay was not successful with the MM253 cells, since they detach easily from the surface, although we used coating method such as collagen.

## Abbreviations

ERK: Extracellular signal-regulated kinase
FBS: Foetal bovine serum
GFP: Green fluorescence protein
IGF1R: Insulin-like growth factor 1 receptor
LIMK: LIM motif containing kinase
OD: Optic density
Ret: Rearranged during transfection
SDS: Sodium dodecyl sulfate
Shc: Src homology collagen
Trk: Tropomyosin receptor kinase
VEGFR: Vascular endothelial growth factor
YPD: Yeast Extract–Peptone–Dextrose

## Contributions

Rayan A. Hago: Performed many of the experiments and participated in writing the manuscript.

Samrein B.M. Ahmed: Supervised the work, conducted some of the experiments, wrote the manuscript and analysed the data.

Sook P. Wong: Screened the mouse embryonic cDNA library by the yeast two hybrid assays.

Mahmood Y. Hachim and Ibrahim Y. Hachim: Helped in analysing patient data obtained from the bc-GenExMiner 4.2 database tool and KM plotter database

Maha Saber-Ayad: Proofread the manuscript and provided valuable comments to enhance the manuscript.

Sally A. Prigent: Provided FLAG-ShcD and GFP-Nischarin constructs, supervised some of the experiments and designed and performed the initial screening

## Data availability

All data generated or analysed during this study are included in this published article.

## Ethics approval and consent to participate

N/A

## Consent to publish

N/A

---

**Figure 1S:**
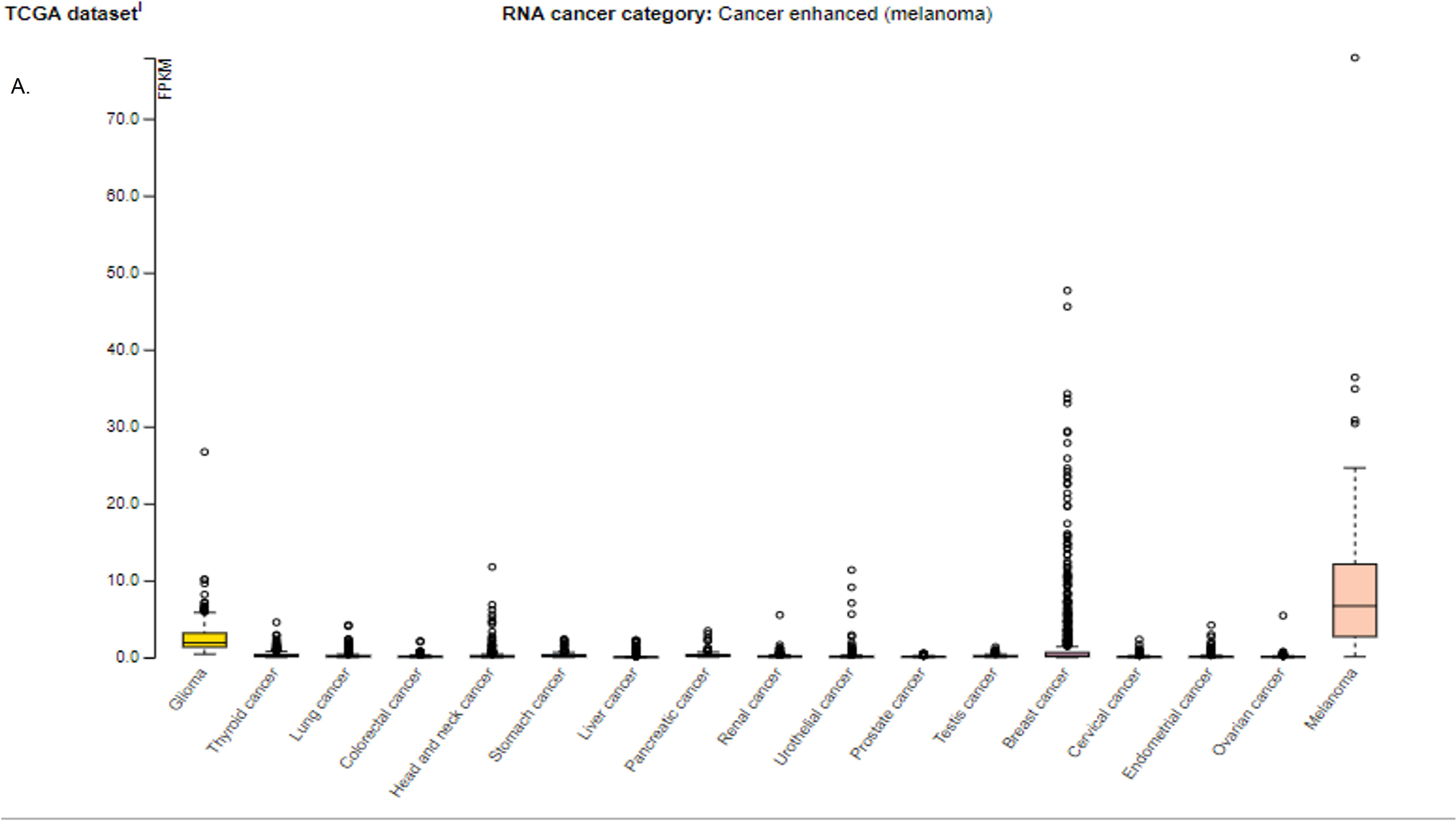
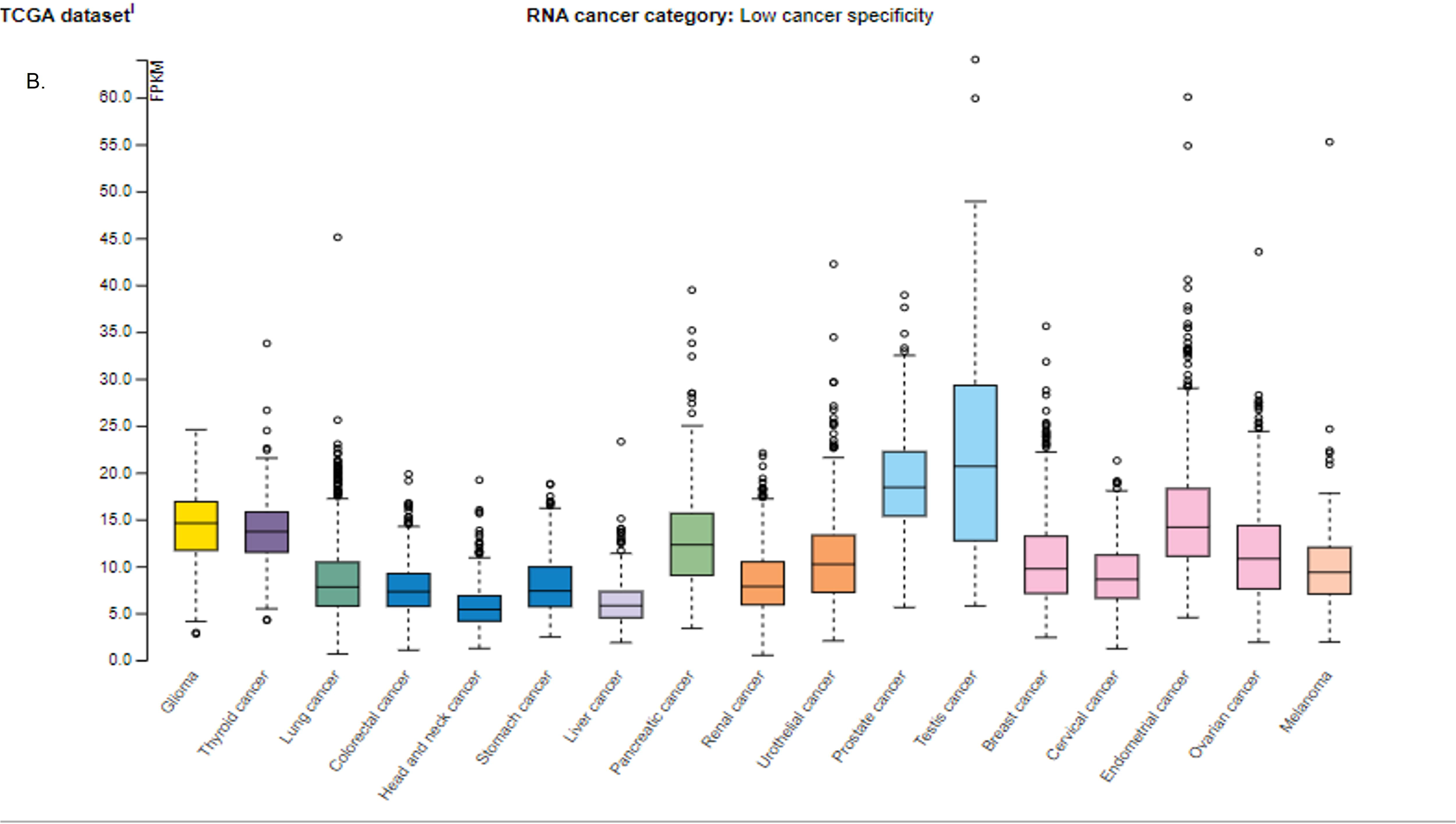
Shc4/D and Nischarin expression in breast cancer and melanoma among other cancers. (A) Shc4/D expression in different cancers, the graph obtained from The Human Protein Atlas at: https://www.proteinatlas.org/ENSG00000185634-SHC4/pathology. (B) Nischarin expression in different cancers, the graph obtained from The Human Protein Atlas at: https://www.proteinatlas.org/ENSG00000010322-NISCH/pathology

**Figure 2S:**
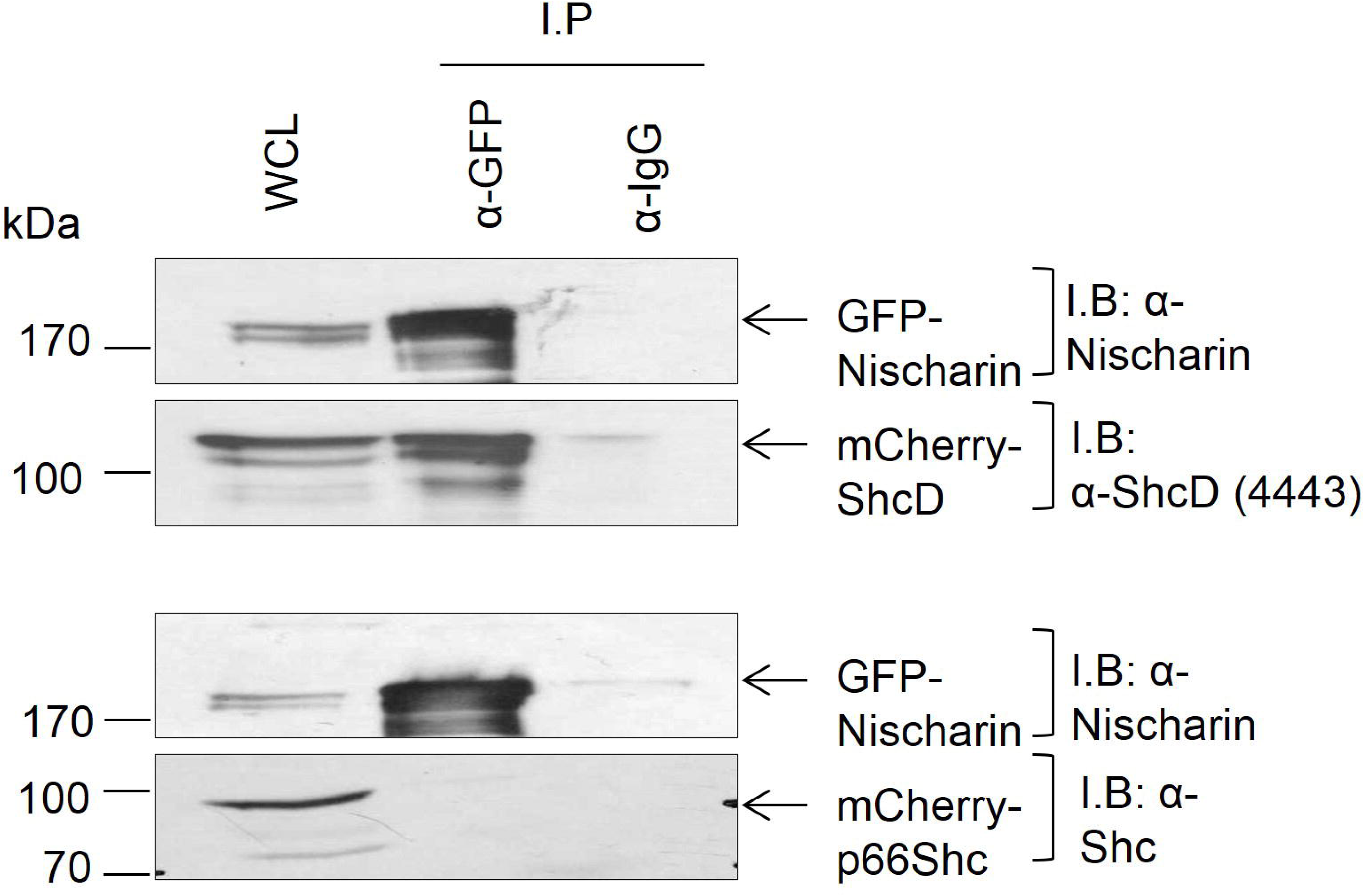
Nischarin binds ShcD but not p66Shc. HEK 293 cells were transfected with GFP-Nischarin and either mCherry-ShcD or mCherry-p66Shc. The cells were lysed, and GFP-Nischarin was immunoprecipitated with 3 µg of anti-GFP antibody. The WCLs represent 1/20 of the total cell lysates. Immunoprecipitates were tested for the presence of mCherry-ShcD or mCherry-p66Shc using anti-ShcD or anti-ShcA antibodies, respectively.

